# *In vivo* imaging of T-cell coregulator B7-H4 reveals protumor macrophage status in prostate cancer

**DOI:** 10.1101/2024.09.28.615608

**Authors:** Manoj Kumar, Shashi B. Singh, Iryna Vasyliv, Frezghi Habte, Mausam Kalita, Israt S. Alam, Sheng Yao Dai, Michelle James, Jianghong Rao, Nicolas Beziere, Heike E. Daldrup-Link

**Affiliations:** Department of Radiology, Molecular Imaging Program at Stanford, Stanford University School of Medicine, Stanford, California 94305, United States; Stanford Cancer Institute, Stanford University, Stanford, California 94305, United States; Department of Neurology and Neurological Sciences, Stanford University, Stanford, California 94305, United States; Department of Chemistry, Stanford University, Stanford, California 94305, United States; Werner Siemens Imaging Center, Department of Preclinical Imaging and Radiopharmacy, Eberhard Karls University of Tübingen, Tübingen, Germany; Cluster of Excellence EXC 2124, “Controlling Microbes to Fight Infections,” Eberhard Karls University of Tübingen, Tübingen, Germany

**Author notes:** Corresponding author: Manoj Kumar, MS, PhD, Molecular Imaging Program at Stanford (MIPS), Department of Radiology, 265 Campus Drive, room G2045, Lorry Lokey Stem Cell Research Building, Stanford University School of Medicine, Stanford, CA 94305-5614.

## Abstract

**Background:** B7-H4 is a cell surface ligand overexpressed by tumors to inhibit T cell functions and evade the immune system. B7-H4 is minimally expressed in normal tissues but is highly expressed by various cancer cells and tumor-associated macrophages (TAM). Despite its importance as an immune checkpoint inhibitor, no imaging techniques specifically targeting B7-H4 have been established. To close this gap, we sought to assess the ability of a monoclonal antibody (mAb) based immunoPET radiotracer to visualize B7-H4 in human and murine prostate cancer models.

**Methods:** Anti-B7-H4 mAb clone 2H9 was functionally characterized for binding to the human and mouse B7-H4 protein. The antibody was conjugated with chelator p-SCN-Bn-Deferoxamine (DFO) and labeled with radioisotope Zirconium-89 (^89^Zr) to obtain immunoPET tracer ^89^Zr-2H9-mAb. The biolayer interferometry method was used to test the binding kinetics of DFO-2H9-mAb compared to that of parental 2H9 mAb. A group of six athymic nude mice with human DU145 prostate tumor xenograft underwent MicroPET imaging after tail vein injection of ∼150µCi ^89^Zr-2H9-mAb or non-binding ^89^Zr-Isotype-mAb. Next, immunocompetent C57BL/6J mice with TRAMP-C2 tumors each were injected with either PBS (n=8), cold 2H9 mAb (10mg/kg) to block B7-H4 (n=6), or chlodronate liposome (15mg/kg) to cause total macrophage depletion (n=6), followed by ^89^Zr-2H9-mAb MicroPET imaging. An *ex vivo* biodistribution assay was performed after 144 hr post radiotracer injection. Tumor radiotracer binding, quantified as a percentage injected dose per gram (%ID/g), was compared between different experimental groups using two-way ANOVA with Bonferroni or Tukey corrections.

**Results:** Immunoconjugation yielded a 2.59 ± 0.08 chelator-to-antibody ratio, and the binding of DFO conjugated 2H9-mAb was similar to that of parental 2H9 mAb, with unaffected affinity in targeting B7-H4 protein moiety. The radiochemical purity of ^89^Zr-2H9-mAb tracer was yielded >95% with an average specific activity of 5µCi/µg antibody. DU145 tumor xenografts demonstrated significantly stronger radiotracer binding at 24, 48, 72, 96, and 120 hr than the non-binding isotype control group. In TRAMP-C2 tumor xenografts, the radiotracer binding in B7-H4 blocked tumors was significantly lower than in the non-blocked PBS-injected group. Macrophage depletion resulted in a significant decrease in tumor binding compared to the control group. ^89^Zr-2H9-mAb could efficiently distinguish tumors with high sensitivity, showing a high correlation between PET imaging and bio-distribution. More importantly, the immunohistochemistry of the harvested tumor revealed no significant difference between the three groups, as discernible through *in vivo* PET imaging.

**Conclusion:** This study highlights the potential of B7-H4 immunoPET imaging for monitoring immunotherapy response. With the emerging potential of B7-H4 blocking as an immunotherapeutic, immunoPET imaging could be readily expanded to patient stratification and therapy monitoring. B7-H4 imaging could augment our understanding of B7-H4 dynamics in response to various therapeutic interventions in clinical trials. The new B7-H4 immunoPET probe is, in principle, clinically translatable.

## INTRODUCTION

Immune checkpoints are the negative regulators of T cell functions and determining factors between tolerance and immunity [1]. Cancer immunotherapy with immune checkpoint inhibitors (ICIs) blocks immune suppressing signals activating T cells to kill malignant cells [2, 3]. Several monoclonal antibodies (mAb) blocking program death receptors (PD-1) or its ligand (PD-L1), cytotoxic T lymphocyte antigen 4 (CTLA-4), and lymphocyte activation gene 3 (LAG-3) are approved for the treatment of patients with melanoma, lung cancer, lymphoma, and renal carcinoma [4–6]. Although more than a decade has passed since the approval of ICIs, the majority of patients still show limited response, and the overall success rate remains unsatisfactory, ranging from 12% to 60%, depending on the tumor type and therapeutic combination [7, 8]. Immunologically cold tumors, such as prostate cancer, inherently exploiting multiple immune suppressive mechanisms, have been challenging to treat with ICI [9–11]. Therefore, understanding the immune factors shaping the antitumor immune response and developing new strategies to improve T-cell function is critical for advancing personalized treatment. The lack of static biomarkers presents substantial challenges due to patients’ heterogeneity and inter- and intra-tumor differences across different tissue sites [12]. Novel non-invasive and dynamic biomarkers could be critical for identifying ICI response, patient-tailored predictions, and personalized treatments. From this perspective, new immunoPET imaging biomarkers could be vital in effectively implementing immunotherapies and immune dynamics [13]. A robust imaging approach that allows for the stratification of patients and the prediction of immunotherapy responses could overcome the limitations of current genomic and immunohistochemical assessment methods by being fast, quantitative, noninvasive, and longitudinal, addressing a pressing clinical need.

The B7 family of immune checkpoints plays crucial roles in regulating T-cell response by providing costimulatory and coinhibitory signals [14]. Among these, B7 homolog 4 (B7-H4) has been shown to play a pivotal role in immunologically cold tumors, growth, and metastasis [15]. B7-H4 encodes a heavily glycosylated protein and inhibits T-cell functions by limiting proliferation, cytokine production, and cytotoxic activity [16–18]. Like its sibling PD-L1 (B7-H1), B7-H4 negatively regulates T-cell-mediated immune responses and is critical in the body’s immune checkpoint mechanisms [19]. While B7-H4 in normal tissues alters the balance between inflammation and tolerance by suppressing T-cells, it is highly expressed across human cancers and protumor macrophages [17, 20, 21]. Tumor-associated macrophages (TAM) also express B7- H4 [22], promoting immune suppressive functions and creating an environment conducive to tumor growth and survival [23, 24]. In the tumor immune microenvironment (TIME), macrophages (Mɸ) are highly polarized towards protumor wound-healing phenotype (M2), which provides a significant source of immunosuppressive signals and blocks T-cell infiltration [25–27]. TAM accounts for about 30-50% of the tumor stromal cell population and has been shown to reduce therapeutic responses and worsen patient outcomes [28]. While several malignancies have shown an increased expression of B7-H4, its presence in prostate cancer is particularly noteworthy [23, 29], as studies suggest, B7-H4 expression correlates with more aggressive prostate cancer phenotypes, higher Gleason scores, and advanced stages [30]. B7-H4 in prostate cancer could directly contribute to the aggressive stage, driving metastasis and potentially mediating resistance to conventional therapeutic modalities [29]. Preclinical studies investigating B7-H4 targeted therapies have shown promising results when utilizing mAb to block B7-H4 as an immune checkpoint [31] or delivering cytotoxic drugs via antibody-drug conjugates [32, 33]. Noninvasive monitoring of B7-H4 status promises to provide valuable insight into tumor immune status, TAM status, and therapy response.

Quantitative imaging of B7-H4 can provide insights into anti-tumor immune mechanisms and could guide the customization of combination therapies to maximize clinical benefits. The lack of clinically translatable imaging tracers for B7-H4 and promising results from recent immunotherapeutic preclinical studies [34] motivated us to develop an immunoPET tracer for *in vivo* imaging of B7-H4. Here, we show that B7-H4 immunoPET imaging enables *in vivo* tracking of B7-H4 protein, primarily expressed in tumors and TAM. We achieved this by labeling an anti- B7-H4 monoclonal antibody (clone 2H9; further referred to as 2H9 mAb) recognizing human and mouse B7-H4 protein, highly conserved, with 91% sequence homology between both species [35–37]. The 2H9 mAb was radiolabeled with ^89^Zr radioisotope using chelator Deferoxamine (DFO). We hypothesized that tumor binding of the B7-H4 immunoPET radiotracer would correlate with *ex vivo* staining of B7-H4 protein and variation in response to treatments affecting B7-H4 expression *in vivo*. Recognizing that tumor-associated macrophages (TAMs) also express B7-H4, we aimed to directly assess the dynamics of radiotracer binding in response to total Mɸ depletion in mice. A mouse cohort was injected with clodronate liposome to deplete Mɸ *in vivo* and was longitudinally imaged using ^89^Zr-2H9-mAb. We present the validation of the radiotracer ^89^Zr-2H9- mAb in human and mouse prostate cancer tumor xenografts, assessing their sensitivity and specificity. To our knowledge, this is the first immunoPET study quantitatively assessing B7-H4 status in the TIME. Our approach could help better understand immunotherapy and guide response evaluation in preclinical and clinical studies.

## RESULTS

### B7-H4 protein expression in human and murine prostate cancer cells

We used human DU145 and murine TRAMP-C2 prostate cancer cell lines for B7-H4 imaging and first assessed the relative expression of B7-H4 protein level in these cells. The western blot of DU145 cell lysate showed a B7-H4 protein band at a higher molecular weight than the TRAMP- C2 cell lysate (Figure 1). The glycosylated forms of B7-H4 were observed as a band of varying size ranges. The non-glycosylated B7-H4 protein is observed as a 28kDa band [38]. A quantitative assessment of western blot bands showed that DU145 cells had a 2.8 ± 0.71 fold higher B7-H4 expression than TRAMP-C2 cells. Further immunofluorescence staining of B7-H4 in both cell lines visually confirms B7-H4 expression (Figure 1B). Thus, these results demonstrate that 2H9 mAb, as detected by anti-mouse secondary antibodies, binds to both human and mouse B7-H4 proteins *in vitro*.

**Figure 1:**
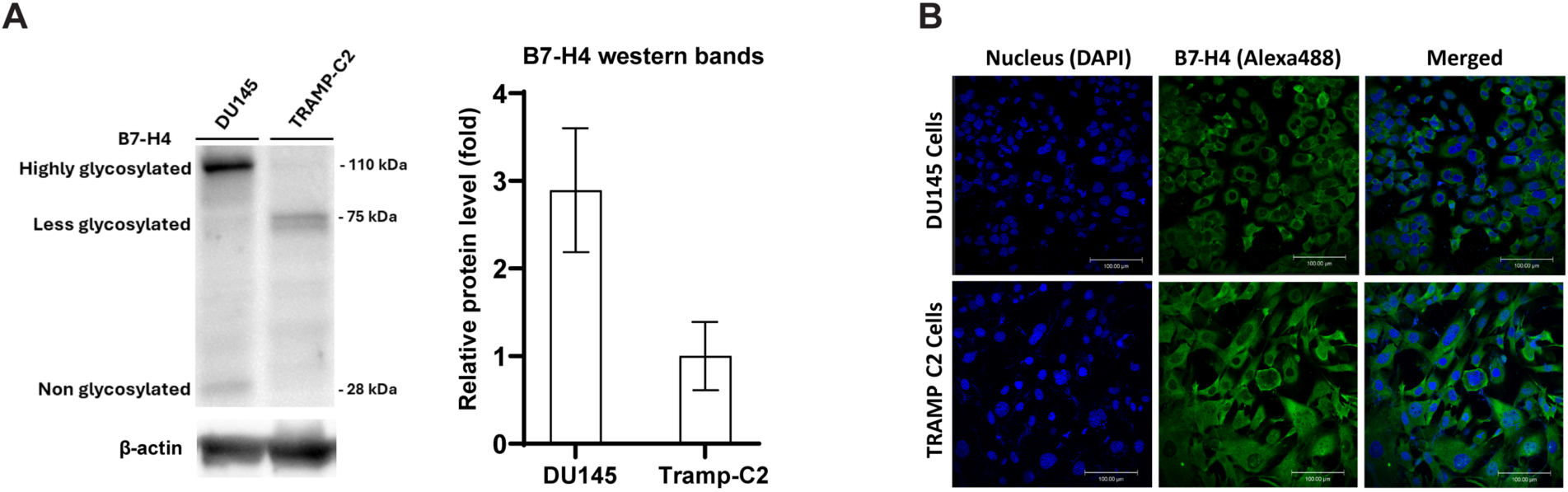
B7-H4 expression in prostate cancer cell lines. **A)** Western blot protein bands showing the expression level of B7-H4 protein in human DU145 and murine TRAMP-C2 cell lines. The bar graph shows the relative expression of B7-H4 protein levels between DU145 cell and TRAMP-C2 cell calculated as fold differences using ImageJ software. Data are displayed as mean data of n = 3 per group and standard deviation. **B)** Immunofluorescence staining of B7-H4 protein in prostate cancer cell lines with DAPI (blue) nuclear staining and B7-H4 (green, Alexa-488). Scale bar = 100µm.

### Development and characterization of anti-B7-H4 immunoPET tracer

Next, we conjugated the 2H9 mAb with chelator DFO to facilitate the radiolabeling for PET tracer development. DFO is a potent chelator of ^89^Zr and is well-established for radiolabeling antibodies in preclinical and clinical studies [39]. The mass spectrometry analysis of the chelate number showed an average of 2.59 ± 0.08 DFO per antibody (Figure 2A). The integrity of the resulting DFO conjugated 2H9 mAb was maintained as reflected by HPSEC/MS peak analysis of modified compared to unmodified parental 2H9 mAb. Subsequently, we tested whether DFO conjugation impacted the affinity of 2H9 mAb. The BLI analysis of modified DFO-2H9 mAb displayed nearly unaffected binding affinity to human and mouse recombinant B7-H4 (rB7-H4) protein compared to binding of unmodified parental 2H9 mAb. When tested with human rB7-H4, the nanomolar affinity constants (K_D_) of DFO-2H9-mAb were measured as 26 ± 1.67 nM compared to 19 ± 0.97 nM of parental 2H9 mAb. Likewise, when tested with mouse rB7-H4, the K_D_ values were 18 ± 0.87 nM for DFO-2H9-mAb compared to 21 ± 1.23 nM for parental 2H9 mAb (Figure 2B). Thus, the obtained BLI binding curves demonstrated unaffected affinity of DFO-2H9-mAb in targeting the B7-H4 protein in both species. Subsequently, the DFO-2H9 mAb was radiolabeled with ^89^Zr, and the radiochemical purity of the resulting radiotracer ^89^Zr-2H9-mAb exceeded 95% in all radiosynthesis (Figure 2C). HPSEC analysis showed no evidence of aggregation (Figure 2D). These results revealed pure ^89^Zr-2H9-mAb radiotracer without any free ^89^Zr in the sample. The elution time of the radioactive peak from ^89^Zr (9.88 min) and the UV peak from 2H9 mAb (9.76 min) align in the size exclusion chromatography, demonstrating intact ^89^Zr-2H9-mAb radiotracer.

**Figure 2:**
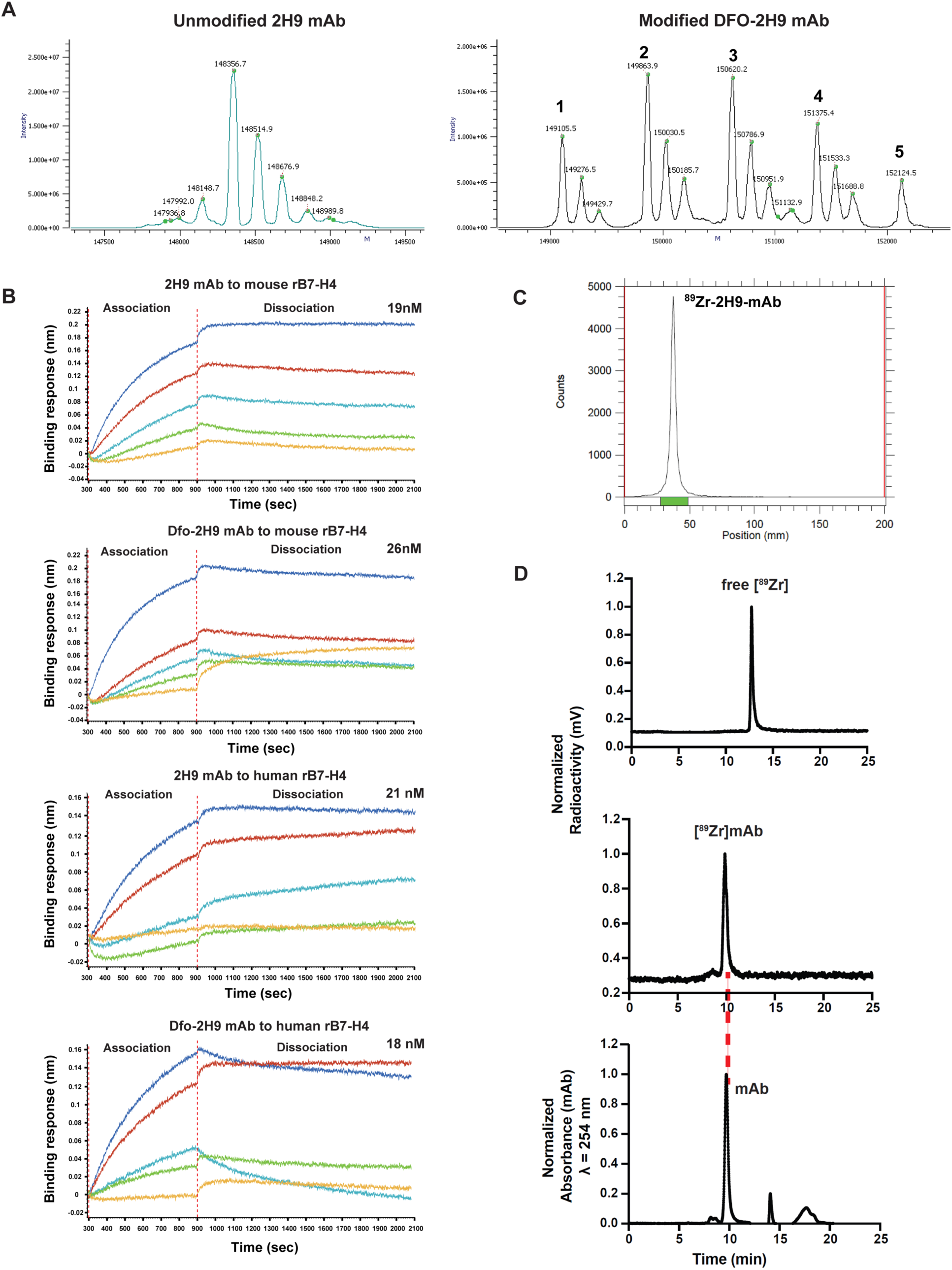
Immunoconjugate, affinity, and radiotracer yield: **A)** SEC-MS characterization of parental and DFO conjugated 2H9 mAb. The mass peaks represent heterogeneity in antibody fragments and molecular weight changes in respective peaks after DFO conjugation. Numbers at the top of the peaks indicate the linkers conjugated per antibody. The glycosylated forms of parental 2H9 antibody fragments ranged from 147936-148989 Da, and the after DFO conjugated antibody size ranged from 149105-152124 Da. **B)** Association and dissociation curves of the parental and DFO conjugated 2H9 mAb to mouse and human recombinant B7-H4 proteins were determined using real-time BLI analysis. The curves obtained were used to determine the affinity (nM) of unmodified and modified 2H9 mAb. **C)** Radio iTLC peak represents the quantitative radiolabeling yield of the final tracer. D) HPLC chromatogram of 89Zr-2H9-mAb.

### 89Zr-2H9-mAb immunePET imaging in mice bearing DU145 xenografts

To test whether we can detect B7-H4 expression *in vivo*, we first established DU145 tumor xenografts in male athymic nude (nu/nu). The average tumor size before radiotracer injection was 134 ± 72 mm^3^. Mice were imaged at 4, 24, 48, 72, 96, and 120 hr post radiotracer injection (Figure 3A), and the tumor binding of the radiotracer at each timepoint was measured as 3.52 ± 0.79, 4.72 ± 0.92, 5.37 ± 0.76, 5.25 ± 0.98, 5.34 ± 1.12, and 5.49 ± 1.08 %ID/g, respectively (Figure 3B).

**Figure 3:**
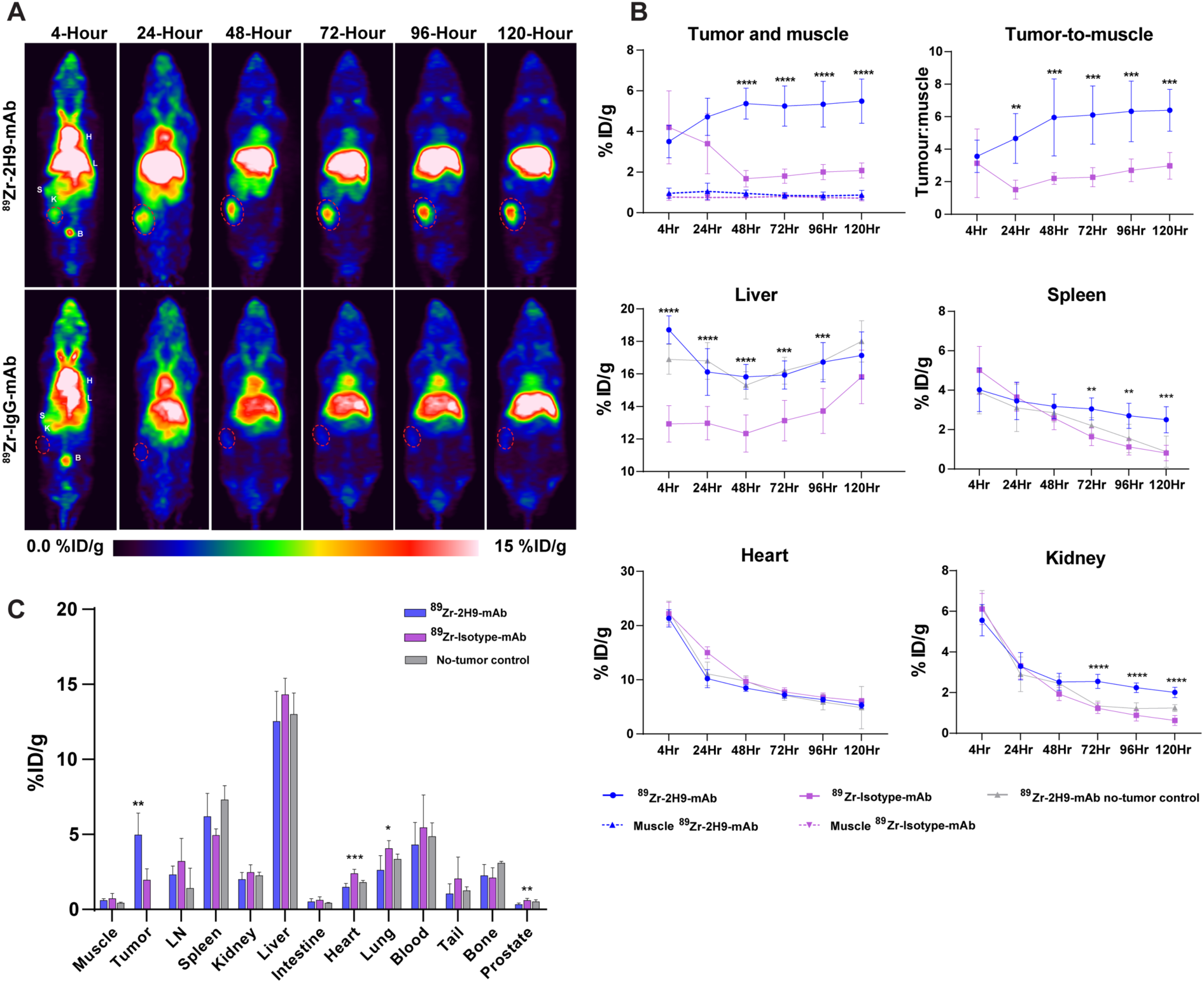
Whole-body PET imaging of athymic nude mice bearing DU145 xenografts. **A)** Maximum intensity projections (MIP) of representative mice from 89Zr-2H9-mAb (top row) and 89Zr-IgG-mAb (bottom row) cohorts at 4 hr to 120 hr imaging time points (n = 6 each). Tumor is circled with red dotted line. **B)** Relative ROI measure of radioactivity (%ID/g) in tumor, muscle, tumor-to-muscle ratio (T:M), liver, heart, spleen, and kidney at each imaging timepoints after intravenous injection of 150µCi activity of radiotracer. **C)** Ex vivo biodistribution of 89Zr-2H9-mAb and 89Zr-Isotype-mAb in tumor and different visceral organs of mice harvested at 144 hr post radiotracer injection (n = 6, no tumor control n = 2). Data are displayed as mean ± standard deviation. H = heart, L = liver, S = spleen, K = kidney, B = bladder, LN = lymph node. *\* = P< 0.05, ** = p<0.01, ***=P<0.001, and P ****=< 0.0001*.

The blood pool activity (ROI on heart) was highest at 4 hr diminished over time, while tumors could be visualized with significantly higher binding at 24 hr post radiotracer injection to the final imaging time point at 120 hr. To demonstrate the specificity of our radiotracer, we injected ^89^Zr-Isotype-mAb (mouse IgG1) as a non-binding control in a second mouse cohort with an average tumor volume of 119 ± 53 mm^3^. The mice cohort injected with ^89^Zr-Isotype-mAb showed no significant tumor tracer binding compared to the ^89^Zr-2H9-mAb cohort. Expectedly, the non-specific tumor binding of ^89^Zr-Isotype-mAb was greater than muscle binding, likely due to enhanced permeability and retention effects in the tumor stroma. When statistically tested, a significant difference in the tumor binding between the ^89^Zr-2H9-mAb and ^89^Zr-Isotype-mAb cohort was observed from 48-hr onwards. Thus, the gradual increase in the tumor binding activity exceeding that of the blood pool and muscle suggests B7-H4 receptor-mediated radiotracer binding and not a non-specific binding due to the enhanced permeability and retention effect. The tumor-to-muscle contrast ratio increased longitudinally and reached 6.39 ± 1.20 at 120 hours post radiotracer injection (Figure 3B). The liver binding was highest among all tissues, which is expected due to the metabolic clearance of radiotracer as a large antibody-based molecule. However, the liver accumulation of ^89^Zr-Isotype-mAb was significantly lower than ^89^Zr-2H9-mAb at all imaging intervals except that of 120 hr imaging (Figure 3B). Likewise, ^89^Zr-2H9-mAb accumulation dropped moderately in the spleen, while ^89^Zr-Isotype-mAb dropped significantly after 72 hr imaging timepoint (Figure 3B). Additionally, we injected ^89^Zr-2H9-mAb in healthy mice with no tumors (N=2) and observed similar organ-specific tracer accumulation compared to mice bearing DU145 tumor xenograft (Figure 3B and C).

The *ex vivo* biodistribution of ^89^Zr-2H9-mAb and ^89^Zr-Isotype-mAb in both cohorts was obtained by measuring radioactive signals in the harvested tissues at 144 hr (Figure 3C). Tumor binding was measured as 4.97 ± 1.45 %ID/g in the ^89^Zr-2H9-mAb cohort compared to 1.96 ± 0.75 in the ^89^Zr-Isotype-mAb cohort. Likewise, the average tumor-to-muscle ratios for the ^89^Zr-2H9-mAb cohort were 8.24 ± 2.14 %ID/g compared to 2.89 ± 1.44 %ID/g in the ^89^Zr-Isotype-mAb cohort. These results indicate that ^89^Zr-2H9-mAb can efficiently detect and differentiate B7-H4- expressing prostate tumors from normal tissues.

### B7-H4 immunoPET imaging reveals tumor-associated macrophage status in syngeneic TRAMP-C2 tumors

Next, we sought to test the utility of ^89^Zr-2H9-mAb PET tracer in an immunocompetent model using TRAMP-C2 tumor xenografts established in male C57BL/6J mice. We asked whether this radiotracer could differentiate the level of B7-H4 expressed by TAM, maintaining high *in vivo* specificity to low and high B7-H4 protein levels *in vivo*. To differentiate the B7-H4 expressed by cancer cells and TAMs, we randomized TRAMP-C2 tumor-bearing mice in three imaging cohorts: the tumor-bearing control as a non-blocked group, the blocked group injected with cold parental 2H9 mAb, and a Mɸ-depleted group injected with clodronate liposome (Figure 4A) [40]. The average tumor size measured was 142 ± 64 mm^3^ before randomizing mice in three groups. All mice were injected with 150*µ*Ci radiotracer and imaged at 4, 24, 48, 72, 96, and 120 hr timepoints, with a maintained clodronate liposome regimen (at 48 hr intervals) in the Mɸ depleted group (Figure 4A). Quantitative analysis of normalized PET slices indicates peak tumor binding signals from 48 hr to 120 hr after tracer injection (Figure 4B, C). The mean tumor binding in each group at each imaging timepoint was measured as 6.26 ± 2.47, 8.48 ± 2.076, 11.99 ± 1.594, 12.40 ± 1.5, 12.60 ± 1.47, and 12.95 ± 1.57 %ID/g in the non-blocked control group (Figure 4B, C, top row), 2.49 ± 1.06, 5.64 ± 2.17, 7.36 ± 1.53, 10.36 ± 1.7, 10.40 ± 1.57, and 10.52 ± 1.43 %ID/g in the blocked group (Figure 4B, C, middle row), and 2.95 ± 1.23, 5.64 ± 1.76, 9.19 ± 1.21, 9.15 ± 1.14, 9.43 ± 1.28, and 8.17 ± 1.7 %ID/g in the Mɸ depleted groups (Figure 4B, C, bottom row).

**Figure 4:**
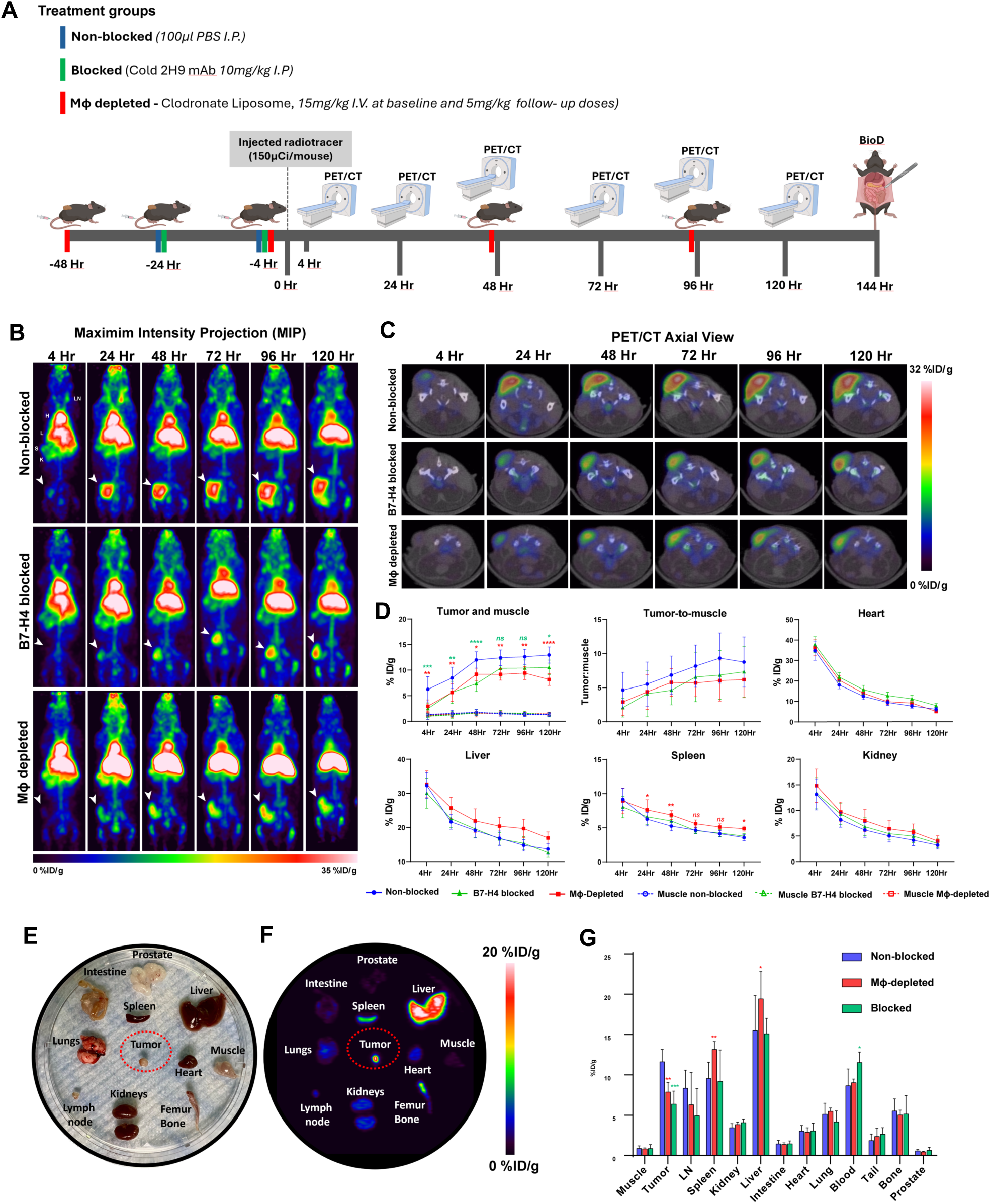
Micro PET imaging of 89Zr-2H9-mAb in TRAMP-C2 tumor bearing immunocompetent mice. **A)** Illustration of PET imaging study design and timeline. Twenty C57BL/J6 mice with subcutaneous TRAMP-C2 tumors were randomized in three imaging cohorts as non-blocked (n=8), blocked (n=6), and Mɸ-depleted (n=6) for longitudinal microPET imaging. All animals were sacrificed at 144 hr after intravenous injection of 150μCi radiotracer. Harvested tissues and organs were loaded on gamma counter for ex vivo biodistribution assay. **B)** Maximum intensity projections (MIP) images of representative mice from each cohort at respective imaging timepoints after radiotracer injection. Tumors are indicated with white arrowhead. **C)** Axial PET/CT slices of the same mouse as in B from non-blocked, blocked, and Mɸ-depleted cohorts. **D)** Percentage of the injected radioactive dose (%ID/g) determined by drawing region of interest in tumor, liver, heart, kidney, spleen and muscle, and calculated tumor-to-muscle ratio (T:M). **E)** Photo of harvested tissues and organs from a representative mouse in non-blocked cohort were arranged on plate **F)** 3D rendered MIP PET image of plate with harvested tissues at mice at 144 hr post radiotracer injection. **G)** %ID/g values of tumor and visceral organs at 144 hr after radiotracer injection, determined from gamma couner readings. H = heart, LN = lymph nodes, L = liver, K = kidney, S = spleen. All data are displayed as mean ± standard deviation. *\* = P< 0.05, ** = p<0.01, ***=P<0.001, and P ****=< 0.0001*. Figure 1A *was created with BioRender*.

Expectedly, the reduced tumor binding in the blocked group verifies the specificity of the B7-H4 blocking effect, showing a significantly reduced tumor PET signal compared to the non-blocked group; the tracer accumulation in these tumors gradually increased over time. More importantly, the tumor binding of radiotracer in the Mɸ depleted mice was significantly lower compared to the non-blocked group (p=0.0020, 0.0090, 0.0102, 0.0026, 0.0030, and 0.001) and remained stable at follow-up timepoints from 48 hr to 120 hr. These observations indicate that the TAM population significantly contributes to B7-H4 PET signals from the tumor stroma, as confirmed by significantly higher signals from the non-blocked control group, and trended lower than the blocked group with no significant difference at any imaging timepoint. Furthermore, the *ex-vivo* PET scanning of harvested tissues from a mouse in the non-blocked cohort identified the accumulation of ^89^Zr-2H9-mAb in the tumor, and signals in other major organs also correlated to the %ID/g values measured *in vivo* via PET imaging (Figure E-F).

Moreover, gamma counter readings quantitatively verified tumor-associated activity as observed in PET imaging of each cohort of mice. The %ID/g radioactivity values determined from gamma counter readings of harvested tissues at 144 hr post radiotracer injection were consistent with those determined by PET ROI analysis at 120 hr timepoint (Figure 4G). While a significant accumulation of ^89^Zr-2H9-mAb was detected in the liver, consistent with the data from DU145 xenografts, we interestingly observed increased tracer accumulation in the liver of the Mɸ depleted cohort when compared to non-blocked and blocked cohorts of mice, these values were statistically not significant. Besides elevated accumulation in the liver, we also observed increased signals in the kidney and spleen of the Mɸ depleted cohort compared to the non-blocked cohort, statistically significant only in the spleen at 24 hr, 48 hr, and 120 hr imaging time points (Figure 4D). This trend was further verified by the gamma counter readings *ex vivo* (Figure 4G), showing elevated gamma counter %ID/g values in the kidney and spleen of Mɸ depleted cohort, and a statistically significant difference was observed only in the spleen (p=0.0049). These results demonstrate that ^89^Zr-2H9-mAb PET signals are specific to B7-H4 and can quantitatively reflect the changing status of B7-H4 and B7-H4 positive TAMs in response to therapy.

### Immunostaining of B7-H4 and total macrophage status in harvested TRAMP-C2 tumors

Next, we utilized the TRAMP-C2 tumor tissues from each cohort of mice, harvested at 144 hr timepoint (Figure 4A), to confirm the level of B7-H4 protein expression and total Mɸ status *ex vivo*. Interestingly, B7-H4 IHC staining between the non-blocked and blocked groups was less apparently different than the difference we observed in PET images of both cohorts (Figure 5A). The B7-H4 staining on tumor sections from the Mɸ-depleted cohort showed marginally faint staining, yet not as apparent as observed through PET imaging. This pattern was also observed in the H&E staining of these tumor sections. Additionally, the western blot of tumor lysates showed a decrease in total B7-H4 protein band intensity from Mɸ depleted tumor lysates (Figure 5B). When protein bands were quantitatively analyzed, this difference was found as statistically not significant.

**Figure 5:**
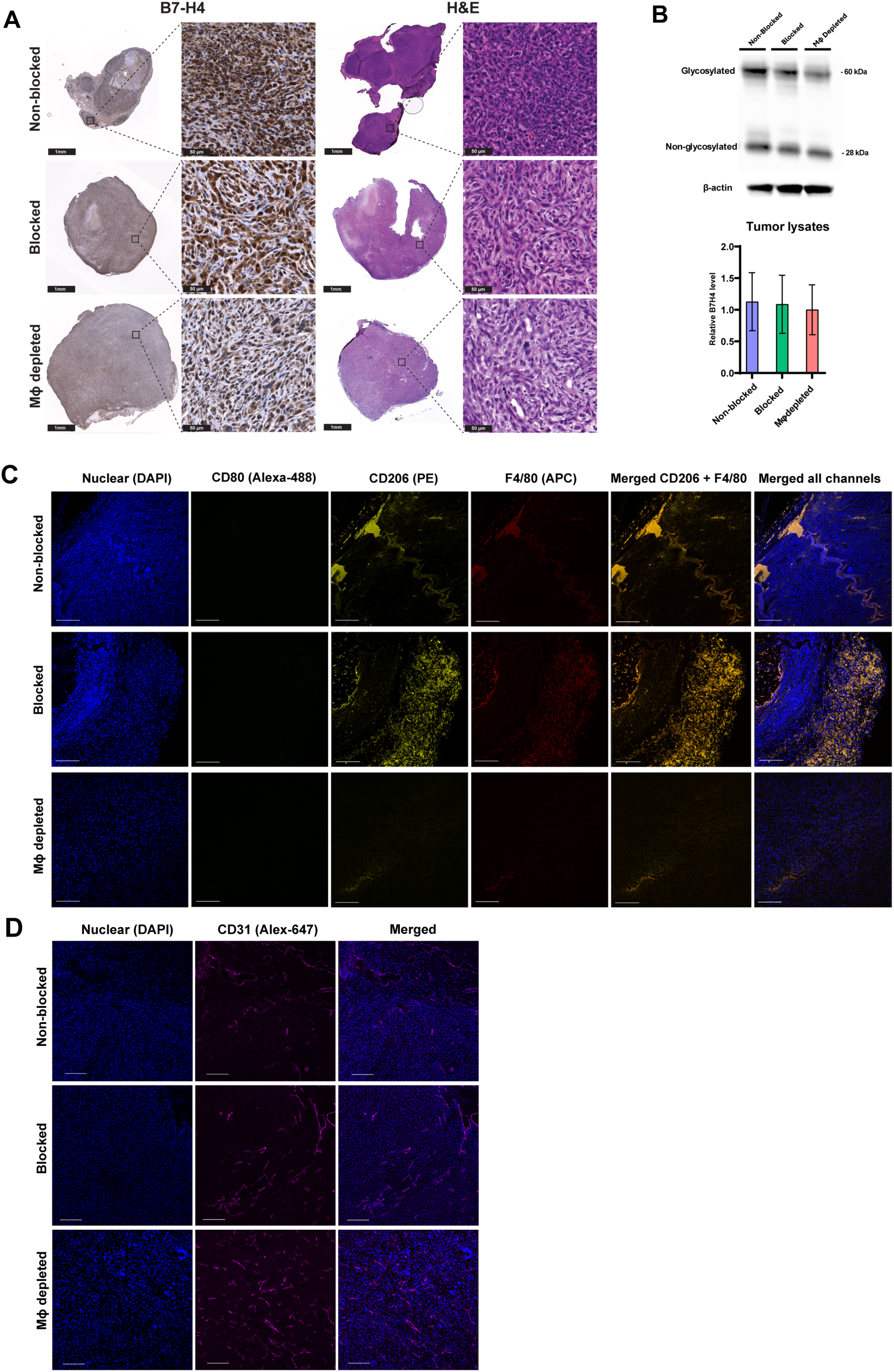
Immunostaining of Tramp-C2 tumors. **A)** B7-H4 protein level in tumors harvested from mice in each imaging cohort stained by IHC method and H&E staining of the same tumors. Whole tumor scale bar = 1mm and magnified image scale bar = 50µm. **B)** Western blot analysis of the tumor lysates from each cohort showing B7-H4 protein bands as glycosylated ∼60kDa and non-glycosylated ∼28kDa bands. The total protein bands in each lane were quantitively analysed using ImageJ software. The bar graph shows the level of B7-H4 protein relative to Mɸ depelted tumor plotted as 1 then the fold differences was calculated for non-blocked and blocked tumors. All values are normalized to β-actin bands from respective lanes. Data are displayed as mean ± standard deviation (n=3). **C)** Multiplexed immunofluorescence staining of the nucleus (DAPI, blue), M1 (CD80, green), M2 (CD206, yellow), and total macrophage (F4/80, red) in the tumor tissue section from each treatment group. **D)** Immunoflourescence staining nuclear (DAPI, blue) and CD31 (magenta) marker of angigenesis in the harvested tumor tissues sections from each cohort. Scale bar = 100µm.

Next, we assessed the Mɸ status on these tumor sections via multiplexed immunofluorescent staining for CD80 (M1 polarized), CD206 (M2 polarized), F4/80 (total), and nuclear DAPI. A high M2-type TAM (CD206) level was detected in the non-blocked control and blocked groups. The tumor sections from the Mɸ depleted group showed a significant decrease in total Mɸ levels compared to non-blocked and blocked groups (Figure 5C). Moreover, extensive blood vessels (CD31 staining) were observed among all three cohorts, showing the active growth stage of all tumors (Figure 5D). These results show that, while B7-H4 levels determined with IHC were not as distinct as quantitatively observed in PET imaging, the total Mɸ status assessed via immunofluorescent staining correlated well with imaging results. The immunofluorescence staining of tumor tissue sections for M1 (CD80) and M2 (CD206) indicate that the vast majority of Mɸ present in the tumor were M2 polarized Mɸ; interestingly, the M1-type Mɸ were rarely observed and were present in only in one tumor tissue sections from the blocked group.

## DISCUSSION

Cancer immunotherapy resistance remains a critical unmet clinical challenge. The development of new imaging probes for investigating the role of potential therapeutic targets, disease diagnosis, patients’ stratification, and effectively monitoring therapy response is of significant interest. There is a critical need for new therapeutic strategies for immune-tolerant tumors to overcome the lack of T-cell infiltration and increase the fraction of patients who can respond to immunotherapy. In this study, we strategically focused on imaging B7-H4, a potent biomarker of cold TIME promoting T-cell inhibition and exhaustion. We utilized an anti-B7-H4 mAb scaffold (clone 2H9) that recognizes common epitopes on human and mouse B7-H4 protein [37]. Using the developed radiotracer ^89^Zr-2H9-mAb, we monitored tumor B7-H4 expression, biodistribution, and immunotherapy response longitudinally in vivo. Like the other B7 family members, B7-H4 is known to be heavily glycosylated [38, 41]. The 2H9 mAb could detect highly glycosylated and non-glycosylated forms of B7-H4 protein bands in prostate cancer cells and tumor lysates. Therefore, we expect that the glycosylation status of the B7-H4 protein does not adversely impact 2H9 mAb binding kinetics.

Further, conjugation of linker DFO to 2H9 mAb enabled radiolabeling by complexation of ^89^Zr. We observed no impact on 2H9 mAb binding affinity to both human and mouse rB7-H4 proteins using the BLI method. The resulting radiotracer ^89^Zr-2H9-mAb was yielded with high radiochemical purity and stability. Longitudinal PET imaging of human DU145 prostate tumor xenografts displayed an excellent tumor-to-background ratio from 48 hr onwards up to 120 hr post-radiotracer injection. PET signal in the tumor resulting from the binding of ^89^Zr-2H9-mAb was well delineated from other tissues. A statistically significant difference in tumor binding of ^89^Zr- 2H9-mAb, compared to ^89^Zr-Isotype-mAb as a reference control, confirmed the high *in vivo* specificity driven by the active binding of the targeting moiety to its B7-H4 epitope. These results show that the radiotracer can differentiate the B7-H4 protein expression level in tumors from healthy tissues. B7-H4 immunoPET imaging represents both tumors and TAMs expressed proteins; therefore, we extended our imaging approach to testing whether ^89^Zr-2H9-mAb can differentiate the *in vivo* status of B7-H4 in response to therapeutic blocking or changes in total Mɸ status *in vivo*. We designed this experimental test using the syngeneic TRAMP-C2 prostate tumors model in immunocompetent C57BL/J6 mice, performing longitudinal immunoPET imaging. Expectedly, B7-H4 blocking with pre-injection of parental 2H9 mAb markedly reduced tumor binding of radiotracer compared to that of the non-blocked cohort, thus reassuring the high specificity of ^89^Zr-2H9-mAb. More importantly, this specificity is also seen in the Mɸ-depleted cohort due to the lack of infiltrating Mɸ in the TIME, which showed a significant decrease in TAM-related tumor binding of the radiotracer compared to the non-blocked control group. Additionally, the *ex vivo* biodistribution analysis showed that the tumor binding of ^89^Zr-2H9-mAb and its overall kinetics measures in major organs were in agreement with that observed from immunoPET imaging. Thus, these results confirmed that TAMs are the significant contributors of B7-H4 signals in tumors, consistent with previous preclinical and clinical studies investigating TAM status in solid cancers [22, 42]. Thus, we show that B7-H4 immunoPET imaging could be a potential marker of response to TAM-targeted immunotherapies and monitoring TAM status.

While imaging with whole mAb-based radiotracers is typically performed 3 to 5 days after administration to achieve an acceptable signal-to-noise ratio, the tumor-to-muscle ratio in our study suggests the optimal time in the syngeneic prostate tumor model was between 48 hr and 120 hr. *Gitto, S.B. et al.* reported B7-H4 expression to be stable when characterized and serially matched in individual patients and increases after chemotherapy treatment [32]. The extended tumor retention of ^89^Zr-2H9-mAb in our preclinical models supports its utility in longitudinal monitoring of B7-H4 when assessing TAM-targeted therapies, especially where biopsy is not feasible from multiple metastatic sites. IHC has limitations for longitudinal monitoring; its semi-quantitative nature could give rise to discordance of results, where the difference in subsequent tissue processing and interpretation often leads to variability, as they represent a single time point and a single spatial assessment [43]. Our results demonstrate that B7-H4 immunoPET imaging could overcome this limitation and could be extended to other cancer types.

Immunostaining of B7-H4 in the harvested tumor tissues was consistent with our observations of ^89^Zr-2H9-mAb tumor signals, where, we observed a decreased staining density in the Mɸ-depleted tumors compared to the tumor sections from the non-blocked and blocked cohort. Considering that these staining differences could result from infiltrating Mɸ in tumor stroma, we further confirmed the status of M1, M2, and total Mɸ via multiplexed immunofluorescence staining of tumor sections from each cohort. The acquired microscopic images revealed a strong presence of M2 polarized Mɸ in the non-blocked control and blocked cohort compared to the Mɸ-depleted cohort. Therefore, based on the changing dynamics of B7-H4 levels we observed in our imaging study, our data show how ^89^Zr-2H9-mAb can reveal dynamic changes in TAM levels in response to Mɸ depletion mediated by liposome clodronate. These results show that immunoPET imaging could show differences in B7-H4 leaves more robustly than IHC staining with real-time measures of therapy response. In efforts to develop a greater understanding of tumor immune biology, essentially to the development of new immunotherapies, B7-H4 PET imaging can identify whether this pathway is active in individual patients and could allow a predictive approach for effectively tailoring combination therapies.

Our study design has several limitations, and our observations warrant further investigations to correlate the B7-H4 level with other systemic factors. First, our study design did not account for the level of other immune cell population that are reported to express B7-H4 [19]. Therefore, these changes in immune dynamics and the composition of different immune cell types can be addressed with appropriate study designs involving the broader assessment of the immune cell population and B7-H4 expression. As observed in our B7-H4 PET imaging, changes in cell populations may also apply to the liver, spleen, and other organs. Liver as a significant site of IgG catabolism in mice, and like in other studies, intense intense signal retention over time. Kupffer cells are liver- resident Mɸ localized within the liver lumen and adherent to endothelial cells; while we may assume that the higher signal in the liver could be partially due to these Kupffer cells and other cells, such as dendritic cells, the endogenous level of B7-H4 protein expression in Kupffer cells is still not well established. We did not detect appreciable variation in total Mɸ level (F4/80) in liver sections from each treatment group [44], which could be investigated in detail in future preclinical/clinical studies. Second, soluble forms of B7-H4 protein have been reported in preclinical and clinical studies [45, 46], and we expect PET imaging of B7-H4 to vary with the level of soluble B7-H4 in individual bloodstreams in advanced disease state. In our research, smaller TRAMP-C2 tumor xenografts showed a clean tumor periphery when compared to leaky signals in larger tumor xenografts at later timepoints, possibly due to soluble B7-H4 accumulation overtime, in the larger volume tumors. Future evaluation of B7-H4 levels in the bloodstream and correlation with PET image analysis could better explain this phenomenon. Third, the radioisotope ^89^Zr is known to display an affinity for phosphate and accumulates in bone. This well-established effect was also observed in our study, which explains the apparent accumulation of signal in the shoulder and knees of mice. However, these sites could additionally be the site of Mɸ-associated B7-H4 signals. The insights we gained from our B7-H4 imaging study design confirm the potential of monitoring B7-H4 status as an immunotherapy target. B7-H4 role in prostate cancer seems intricately involved in the disease’s pathobiology. As research on B7-H4 targeted therapies deepens, the hope is that a clearer understanding of B7-H4 biology will pave the way for more effective, tailored immunotherapy for cancer patients.

In conclusion, B7-H4 radiotracer ^89^Zr-2H9-mAb showed preferential tumor accumulation. B7-H4 blocking *in vivo* and Mɸ depletion reduce the tumor accumulation of the new radiotracer. We successfully designed a B7-H4 imaging immunoPET tracer and showed that TRAMP-C2 prostate tumors significantly attracted a TAM population expressing B7-H4. This study represents the first dynamic imaging of B7-H4 expression *in vivo*. Our results are sufficiently promising to warrant further investigation on understanding the B7-H4 dynamics through PET imaging.

## METHODS

### Cell culture

Human and mouse prostate cancer cell lines DU145 and TRAMP-C2 were purchased from the American Type Culture Collection (ATCC). Cells were maintained in RPMI 1640 medium with 1% penicillin/streptomycin and 10% fetal bovine serum (FBS) at 37°C and 5% CO_2_. Cells were routinely tested for mycoplasma using the MycoAlert^TM^ Mycoplasma Detection Kit (LONZA).

### Immunoblotting

DU145 and TRAMP-C2 cells were washed with cold PBS and scraped off with RIPA lysis buffer supplemented with protease and phosphatase inhibitors. Cells were lysed on ice and centrifuged at 12,000 rpm for 15 minutes at 4^0^C. Total protein concentrations were detected via the BCA protein assay kit, and samples in the SDS sample buffer were heat-denatured at 95°C for 10 min. Sample proteins (50µg) were loaded and separated by SDS PAGE and transferred into a nitrocellulose membrane. The bands of B7-H4 protein were blotted with primary antibodies, including anti-B7- H4 (2μg/mL dilution, clone 2H9, Creative Biolabs, NY, USA), anti-β-actin (1:10,000, Abcam) and then incubated with secondary antibody goat anti-mouse IgG-HRP (Abcam, Ab97240). Bands were detected using BioRad^TM^ ECL western blot substrate and scanned on BioRad GelDoc XR+. The relative fold of protein bands was quantified using ImageJ software. The values obtained were normalized to β-actin levels in respective lanes.

### Immunofluorescence staining

DU145 and TRAMP-C2 cells were cultured in a 35mm microscopy dish for 48 hours before fixation and washed for staining. Cells were treated via the general immunofluorescence staining procedure. Briefly, cells were fixed and washed with 4% paraformaldehyde (PFA) and PBS. After blockage with 2% goat serum, samples were incubated with the anti-B7-H4 2H9 mAb with a final concentration of 10μg/mL at 4^0^C overnight. 1µg/mL dilution of Goat anti-Mouse IgG (H+L) Cross-Adsorbed Secondary Antibody, Alexa Fluor™ 488 (Thermo Fisher Scientific) was applied to develop immunofluorescence and slides were mounted with ProLong™ Gold Antifade Mountant with DNA Stain DAPI (Thermo Fisher Scientific). The immunofluorescence images of cells were obtained on a DMi8 confocal microscope (Leica Microsystems).

### Radioimmunoconjugation

The conjugation of linker deferoxamine (p-SCN-Bn-DFO, Macrocyclics, USA) and ^89^Zr labeling of 2H9 mAb were conducted using the same strategies reported previously [39]. In brief, antagonist therapeutic anti-B7-H4 mAb (mouse IgG1, clone 2H9, Creative Biolabs, NY, USA) was buffer exchanged with PBS, and pH was adjusted to 8.5-9.0 using 0.2 M Na_2_CO_3._ Next, chelator p-SCN-Bn-DFO in dimethyl sulfoxide (DMSO) was mixed at a molar ratio of 1:15, followed by 1-hour incubation at 37° C on a tube shaker at 400 rpm. The resulting product was purified via a PD-10 desalting cartridge. The antibody-to-DFO ratio was determined by peaks obtained from Electrospray Ionization Mass Spectrometry (ESI-MS) analysis performed for unmodified parental and DFO-conjugate antibodies. The number of DFO molecules per antibody was determined by dividing the m/s ratio difference between the peaks of the whole antibody conjugated and unconjugated with DFO molecules (752.9 Da).

^89^Zr was purchased from a PET trace cyclotron (3D imaging, Little Rock, AR) for radiochemistry. ^89^Zr-oxalate (37 MBq) was added to 0.5 mM HEPES buffer (pH 7.0) and incubated with DFO-conjugated antibodies (∼ 200 µg) at 37 °C for 60 min. Incorporation of the radioisotope was confirmed via instant iTLC analysis (≥99% radiochemical purity (RCP); iTLC-SG (Agilent), mobile phase 10 mM EDTA). The obtained radiolabeling yield and radiochemical purity were routinely >95%, and no purification step was performed after the final yield. The tracer ^89^Zr-2H9- mAb was formulated in 0.9% NaCl for *in vivo* administration. A non-binding isotype antibody, mouse IgG (clone MOPC-21, BP0083, BioXcell, NH, USA), was conjugated and radiolabeled via the same strategy to obtain ^89^Zr-Isotype-mAb. The identity of the elution profile with the original protein and radiolabeling of the protein peak was confirmed by high-performance size exclusion chromatography (HPSEC) analysis (BioSep SEC s2000, 300×7.8 mm, Phenomenex).

### Biolayer Interferometry analysis

The binding kinetics of the parental anti-2H9 mAb and DFO-conjugated mAb were determined using the Bio-layer interferometry optical system (ForteBio RED384 Label-Free Detection Systems, Sartorius). 2H9 mAb interaction with recombinant human (Acro Biosystems, DE, USA) and mouse B4-H4 (MedChemExpress, NJ, USA) proteins was measured following the optimized Octet User Manual. Octet AMC Biosensors were soaked in ddH2O for 10 min to remove the coating. The assay had six steps: initial baseline (180s), loading (600s), wash (180s), baseline (300s), association (600s), and dissociation (1200s). 2H9 mAb was immobilized on the anti-mouse AMC Biosensors during the loading step with kinetic buffer (PBS with 0.01% Tween 20 and 0.05% BSA) at 80µL final reagent (20µg/mL 2H9 mAb) in the black 384-well microplate to yield a wavelength shift response signal in the range of 0 to 0.2 nm. The concentrations of human and mouse rB7-H4 (100, 50, 25, 12.5, 6.25, 3.125, 1.56, and 0.0 nM) for the association and dissociation steps were prepared in PBS at 80µL final reagent in black 384-well microplate. The response signal of negative control (no rB7-H4 protein loaded) biosensors in each step was used to subtract experimental values before further data processing. Sensorgrams were generated by Octet Data Analysis HT 10.0.3.7 software, and then processed data values were exported with association and dissociation plots. The equilibrium dissociation constant (K_D_) was calculated sequentially by measuring association and dissociation rates.

### Animal models

All animal studies adhered to Stanford’s Administrative Panel on Laboratory Animal Care (APLAC) guidelines and approved protocol APLAC-12040. Male athymic nude mice and immunocompetent C57BL/6J mice (8–10 weeks) were purchased from Jackson Laboratories, ME, USA. To generate tumor xenografts, 12 athymic nude mice received subcutaneous injection of human DU145 cells in 100µl PBS into the left flank. To generate TRAMP-C2 tumors in C57BL/J6 mice, initially, five mice were subcutaneously injected with three million TRAMP-C2 cells on the left flank. Tumor growth was monitored and measured using calipers, and volume was calculated [(length x width^2^)/2]. After tumors grew above 150mm^3^ (4/5 mice grew tumors), mice were euthanized, and excised tumors were sliced as ∼1mm^3^ tissue chunks and were serially implanted on the left flank of 24 mice. When all palpable tumors were measured at a minimum above 100mm^3^, mice were randomized to three treatment groups before injecting radiotracer. Mice were intravenously administered with either 100*μ*l PBS as vehicle control, parental anti-B7-H4 mAb to block B7-H4 in mice, or chlodronate liposome (Liposome BV, Amsterdam, Netherlands) to deplete total Mɸ in mice. The parental 2H9 mAb was given at 10 mg/kg mouse body weight at 24 hr and 4 hr before radiotracer injection. The *in vivo* depletion of Mɸ in the third cohort was achieved by injection of liposome clodronate starting at 48 hr before radiotracer injection with an initial 15mg/kg dose, then followed up by 5mg/kg subsequent doses every 48 hr up to the final imaging timepoint (Figure 4A). To ensure that Mɸ remained maximally depleted in the tumor. Additionally, mice were subcutaneously injected with 5mg/kg of liposome clodronate near the tumor sites.

### Small animal PET imaging and *ex-vivo* biodistribution

For microPET imaging, mice were anesthetized with 1.5-2 % isoflurane, injected with 150*µ*Ci (5.55 MBq) ^89^Zr-2H9-mAb via tail vein, and scanned in the prone position in a dedicated small-animal PET/CT scanners (The GNEXT PET-CT, SOFIE Biosciences or Siemens Inveon DPET scanner) at 6, 24, 48, 72, 96, and 120 hr intervals post radiotracer injection. After the last PET scan timepoint, all mice were euthanized *via* CO_2_ asphyxiation, and major organs (*e.g.,* heart, liver, spleen, kidney, tibia bone, blood, thigh muscles, and tumor) were harvested for *ex vivo* radiotracer accumulation measurements on a gamma counter (Hidex AMG Automatic Gamma Counter). Before loading tissues into gamma counter tubes for radioactivity measurements, all tissues were measured for wet weight, and radiotracer accumulation was presented as a percentage of the injected dose per gram of organ (%ID/g). Image analysis was performed by co-registering the PET/CT images using Inveon Research Workplace 3.0 (Siemens Medical Solutions Malvern, PA). The region of interest (ROI) was drawn within the tumors, heart, liver, spleen, kidney, and quadriceps as nontarget tissue to analyze the ^89^Zr-Dfo-2H9-mAb pharmacokinetics. The quantitative measurement of PET signals was expressed as the mean percentage injected dose per gram of body weight (%ID/g).

### Immunohistochemistry and multiplexed immunofluorescence

Slices of excised tumors and tissue were fixed or flash-frozen for *ex-vivo* analysis. Excised tumors were fixed in 4% paraformaldehyde (PFA), embedded in paraffin, and sectioned (5µm) on slides for staining. Immunostaining was performed following general immunohistochemistry (IHC) and immunofluorescence staining procedures. Briefly, slides were deparaffinized, followed by heat epitope retrieval in citrate buffer (pH 6.0) for 60 min at 95°C. After antigen retrieval, blocking was performed following the protocol using a mouse-on-mouse polymer IHC kit (Abcam). Primary antibodies were added at 1:50 and incubated overnight at 4°C. For HRP-based staining, slides were incubated in DAB chromogen solution for 5 minutes. Immunofluorescence slides were stained with DAPI nuclear stain and mounted after a final wash with TBS buffer. Images were acquired using the NanoZoomer Hamamatsu Slide Scanner or multichannel microscopy using the Leica DMi8 confocal microscope (Leica Microsystems).

### Statistical Analysis

All data were analyzed using PRISM 9 (GraphPad). An unpaired student t-test was conducted for western blot results. PET imaging and bio-distribution data were compared using two-way ANOVA analysis with Bonferroni correction, and Tukey correction was used for multiple tests when comparing more than two groups. Results are presented as the mean ± standard deviation (SD). A p-value of less than 0.05 was considered significant.

## ACKNOWLEDGMENT

Manoj Kumar was supported by the Sanjiv Sam Gambhir-PHILIPS Fellowship program through the Precision Health and Integrated Diagnostics Center at Stanford (PHIND). The authors thank members of the Stanford Center for Innovation in In vivo Imaging (Sci3), and the Cyclotron and Radiochemistry Facility (CRF).

## ABBREVIATIONS

2H9: anti-B7-H4 (Clone 2H9)
^89^Zr-2H9-mAb: anti-B7-H4 Radiotracer
CT: Computed tomography
mAb: Monoclonal antibody
Mɸ: Macrophages
DFO: p-SCN-Bn-Deferoxamine
PET: Positron emission tomography
TAM: Tumor-associated macrophages
TIME: Tumor immune microenvironment

## AUTHORSHIP CONTRIBUTIONS

MKu and HDL conceived and designed the research. MKu, SS, and FH conducted in vivo experiments. MKu, IV, and FH analyzed the data. MKu, MKa, and SYD generated and validated antibody conjugation. MKu and MKa performed the radiolabeling and validation study. MKu performed histological staining and analyzed the data. MKu, IA, NB, JR, MJ, and HDL supervised the study and interpreted the results. MKu, NB, and HDL wrote the manuscript. All authors reviewed and edited the manuscript and approved the final version of the manuscript.

## COMPETING INTERESTS

The authors have declared that no competing interest exists.

## Notes

### Competing Interest Statement

The authors have declared no competing interest.

### Summary of Updates

This version of the manuscript has been revised to remove the redundant figure legend from the body text in the result section 5.

